# Epigenetic small molecule screening identifies a new HDACi compound for ameliorating Duchenne muscular dystrophy

**DOI:** 10.1101/2025.01.24.634796

**Authors:** Ke’ale W. Louie, Eva H. Hasegawa, Gist H. Farr, Amanda Ignacz, Alison Paguio, Alyssa Maenza, Alison G. Paquette, Clarissa Henry, Lisa Maves

## Abstract

Duchenne muscular dystrophy (DMD) is the most common inherited muscle disease. There are currently few effective therapies to treat the disease, although many approaches are being pursued. Certain histone deacetylase inhibitors (HDACi) have been shown to ameliorate DMD phenotypes in mouse and zebrafish animal models, and the HDACi givinostat has recently gained FDA approval for DMD. Our goal was to identify additional HDACi, or other classes of epigenetic small molecules, that are beneficial for DMD. Using an established animal model for DMD, the zebrafish *dmd* mutant strain *sapje*, we screened a library of over 800 epigenetic small molecules of various classes. We used a quantitative muscle birefringence assay to assess and compare the effects of these small molecule treatments on *dmd* mutant zebrafish skeletal muscle. Our screening identified a new HDACi, SR-4370, that ameliorated *dmd* mutant zebrafish skeletal muscle degeneration, in addition to HDACi previously shown to improve *dmd* zebrafish. We find that a single early treatment of HDACi can ameliorate *dmd* zebrafish. Furthermore, we find that HDACi that improve *dmd* muscle also cause increased histone acetylation in zebrafish larvae, whereas givinostat does not appear to increase histone acetylation or improve zebrafish *dmd* muscle. Our results add to the growing evidence that HDACi are promising candidates for treating DMD. Our study also provides further support for the effectiveness of small-molecule screening in *dmd* zebrafish.

**Graphical abstract:** 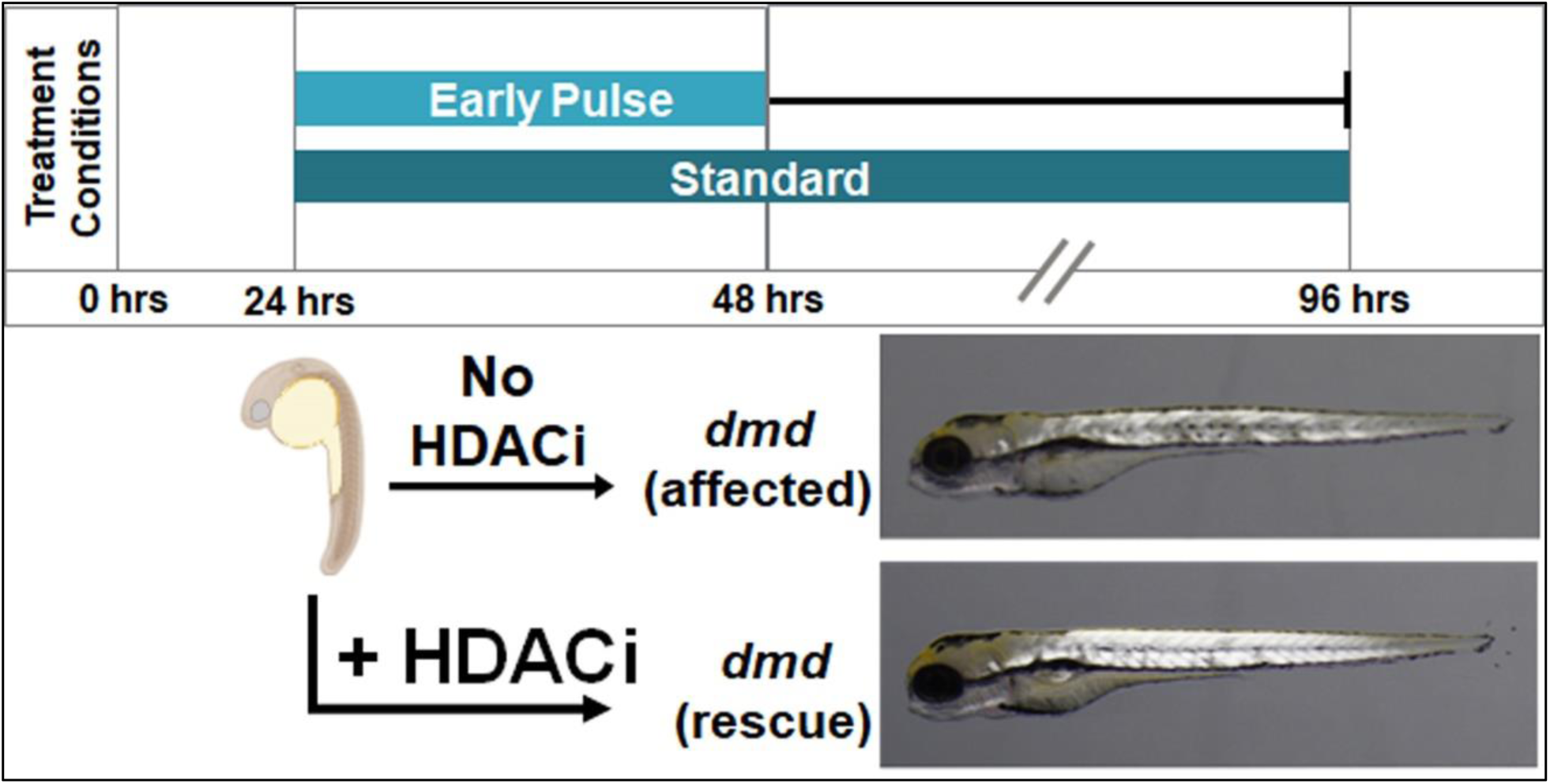

## 1. Introduction

Duchene muscular dystrophy (DMD) is an X-linked muscle degeneration disease caused by mutations in the *DMD* gene, which encodes dystrophin [1], [2], [3]. Dystrophin is associated with the sarcolemma and is part of the dystrophin-associated protein complex (DAPC), which connects the extracellular matrix to the actin cytoskeleton. Mutations in the 2.4 Mb *DMD* gene that result in loss or dysfunction of dystrophin can vary from point mutations to large deletions [3], [4]. Loss of dystrophin leads to contraction-induced structural myofiber damage and the dysregulation of intracellular signaling pathways [5], [6]. DMD disease progression is characterized by progressive muscle damage and inflammation, fibrosis, and necrosis [3], [7], [8]. Over time, patients become wheelchair-bound due to muscle weakness and eventually experience cardiorespiratory failure. While advances in palliative care have increased the life expectancy of DMD boys from approximately 15 years to the mid-twenties to early forties, there is still no effective cure for the disease.

The current standard of care for DMD is corticosteroid treatment [7], [8], [9]. Corticosteroids such as prednisone, prednisolone, and deflazacort are administered to delay symptoms and ameliorate symptom severity, yet these treatments cause significant adverse side effects [9]. Additional treatment approaches currently under development include using antisense oligonucleotides to induce exon skipping of *DMD* mutations, CRISPR editing, and *DMD* gene therapy [9], [10], [11], [12]. While these approaches are gaining FDA approvals, many issues remain, including efficacy, safety, and delivery [9], [10], [11], [12]. Furthermore, because DMD can be caused by a spectrum of mutations, *DMD* gene-directed therapies that target specific exons or mutations are only applicable to a subset of patients.

There have been significant efforts to develop small molecule therapies that target various disease mechanisms downstream of the loss of dystrophin [13], [14]. Treatment with histone deacetylase inhibitors (HDACi) have shown promising results for DMD in mice and zebrafish animal models as well as in DMD patients [15], [16], [17], [18], [19], [20], [21], [22]. For example, in mice, Minetti et al. [19] showed improved muscle phenotype in *mdx* mice treated with the pan-HDACi trichostatin A (TSA). In zebrafish, both TSA and a combination of oxamflatin and salermide have been shown to reduce the severity of muscle lesions in *dmd* mutant embryos [16], [17]. In a phase 2 and phase 3 clinical trial, treatment with givinostat, a pan-HDACi, improved muscle histology and also improved a stair-climb functional assessment in DMD boys [21], [22]. Recently, the FDA approved the use of givinostat for DMD boys. While HDACi show promise for DMD, the long-term safety and efficacy of HDACi in DMD patients is still being studied [22]. Gaining greater insight into this class of small molecules, including their dosing, safety, and mechanisms of action, could help advance their therapeutic use for DMD [23].

In this study, we tested a commercial library of over 800 epigenetic small molecules for their ability to improve the muscle degeneration phenotype of *dmd* zebrafish, a valuable DMD animal model for drug screening [24], [25], [26], [27]. Zebrafish *dmd* mutants exhibit DMD phenotypes such as skeletal muscle lesions, inflammation, fibrosis, and lethality during the early juvenile phase, making *dmd* zebrafish a well-suited model for DMD [25], [28], [29]. We performed the drug library screen by treating *dmd* zebrafish with compounds pooled from each row and column of each library plate, thereby testing each compound twice. We identify a novel HDACi for DMD, SR-4370, that reproducibly rescues the *dmd* zebrafish muscle lesion phenotype. We also identify optimal doses and timing of HDACi treatments for *dmd* zebrafish. Furthermore, we identify a correlation between HDACi rescue of *dmd* muscle lesions and histone acetylation levels in zebrafish larvae. Our results provide further support for HDACi as beneficial small molecules for DMD and underscore the value of using *dmd* zebrafish for drug screening.

## 2. Materials and methods

### 2.1. Zebrafish husbandry

All animal experiments were carried out in accordance with the Institutional Animal Care and Use Committees (IACUC) at Seattle Children’s Research Institute and at the University of Maine. In general, our approaches followed the recommended standards for zebrafish drug screening [30]. Zebrafish were raised and staged as previously described [31]. Time (hpf or dpf) refers to hours or days post-fertilization at 28.5°C. Eggs were collected from 20–30 min spawning intervals and raised in Petri dishes in ICS water [17] in a dark 28.5°C incubator, up to 5 dpf. After 5 dpf, the fish were maintained on a recirculating water system (Aquaneering, San Diego, CA) under a 14 h on, 10 h off light cycle. From 6–30 dpf, the fish were raised in 2.8 L tanks with a density of no more than 50 fish per tank and were fed a standard diet of paramecia (Carolina) one time per day and Zeigler AP100 dry larval diet two times per day. From 30 dpf onwards, the fish were raised in 6 L tanks with a density of no more than 50 fish per tank and were fed a standard diet of Artemia nauplii (Brine Shrimp Direct) and Zeigler adult zebrafish feed, each two times per day. The wild-type stock and genetic background used was AB. The zebrafish *dmd^ta222a^* mutant line (also known as *sapje*; hereafter referred to as *dmd*) has been previously described and is a recessive, non-sex-linked null allele [24], [25]. *dmd^ta222a^*genotyping was performed as previously described [32]. Experimental zebrafish embryos and larvae were used at stages prior to sexual maturity, and zebrafish lack a definite sex-determining chromosome [33]. Therefore, we cannot account for numbers of experimental animals used from each sex.

### 2.2. Small molecules

Chemical screening was performed using an epigenetic compound library (MedChemExpress, Monmouth Junction, NJ; Catalog #HY-L005) consisting of 817 compounds distributed over fourteen 96-well plates. Most compounds were received as 10 mM stocks pre-dissolved in dimethyl sulfoxide (DMSO). The identity, plate, and well location of each compound is provided in Table S1.

For each plate, compounds across a single row (*i.e.*, rows A-H) or down a single column (*i.e.*, columns 2-11) were combined in equal parts (Fig. 1A), resulting in 185 unique pools. 1000X stocks of chemicals were made in DMSO just prior to each treatment experiment.

**Fig. 1.**
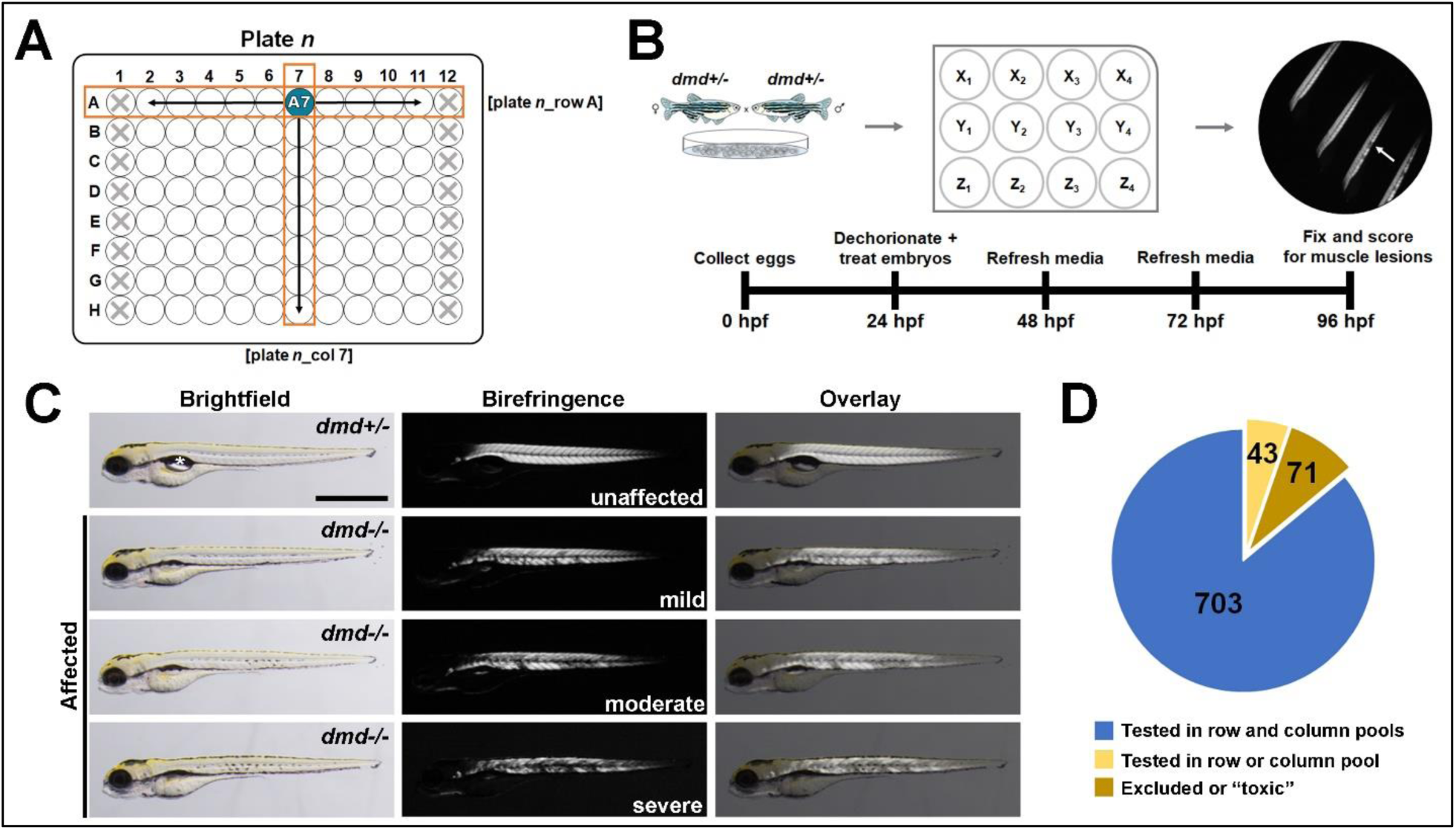
Drug-pool-based epigenetic library screening with *dmd* zebrafish. **A.** Schematic representation of the pooled screening approach. Compounds from each of the individual library plates (n=14) were pooled across rows as well as down columns, yielding 185 unique total pools of 8-10 compounds each. Drug pools were named as [plate n_row x] or [plate n_column y]. Re-tested secondary pools were designated with the suffix “_sec”. Plate columns 1 and 12 were empty across the library (represented as crossed out wells). **B.** Timeline of small molecule treatments. Embryos from *dmd+/−* crosses were collected, dechorionated, and transdermally exposed to compounds or 0.5% DMSO at a density of 25 animals/3 mL media. Media and compounds were replaced daily. At 96 hours post fertilization (hpf), animals were fixed and then scored using polarized light birefringence. Arrow points to muscle lesion. **C.** Brightfield, corresponding birefringence, and overlay images of 96 hpf larvae illustrating representative examples of unaffected and affected animals. Animal genotypes are shown. Affected animals are qualitatively scored as mild, moderate, and severe. Scale bar = 1mm. * = swim bladder. **D.** Distribution of screened library compounds. Over 86% (703 of 817 compounds) were successfully tested in intersecting row and column pools. Compounds at toxic row and column pool intersections were excluded from further analysis (n=71). Compounds in the 5 toxic pools following re-testing (n=43) were only tested in either row or column pools and thus were not assigned composite rescue scores (see Tables S1 and S4).

Select compounds were separately ordered from MedChemExpress or other sources and resuspended in DMSO (Sigma-Aldrich, St. Louis, MO) to make 10 mM stocks. The individually-ordered compounds, sources, and corresponding PubChem CIDs are listed in Table S2.

### 2.3. Small molecule treatments

For epigenetic library screening for *dmd* birefringence rescue tests, compounds were administered from 24-96 hpf as previously described (Fig. 1B) [17], [29]. Embryos from *dmd^ta222a^*^/+^crosses were produced via group spawnings, collected at 20-30 min intervals, sorted to a density of ≤100 embryos/100mm Petri dish, and raised at 28.5°C. Clutches were selected for both appropriate and synchronous development at about 24 hpf as a requirement for experimental inclusion. At about 24 hpf, embryos were enzymatically dechorionated with 0.5 mg/mL Pronase (Sigma-Aldrich, St. Louis, MO) for 14 min and sorted into 12-well plates at a density of 25 embryos/well, as previously described [34]. Quadruplicate wells each received 3 mL of embryo medium (EM; 14.97 mM NaCl, 0.50 mM KCl, 0.98 mM CaCl_2_.2H_2_O, 0.15 mM KH_2_PO_4_, 0.99 mM MgSO_4_.7H2O, 0.05 mM Na_2_HPO_4_, 0.83 mM NaHCO_3_) with either vehicle control or chemical treatment, and treatment media were refreshed every 24 hr. Unless otherwise noted, vehicle control was 0.5% DMSO and the working concentration of compound(s) in a treatment group was 1 µM/compound, based on our previous study [17]. Larvae at 96 hpf were fixed in 10% formalin in phosphate buffered saline (PBS) and stored at 4°C until phenotypic scoring and imaging.

For treatments of individual drugs, we followed the steps described above, except that drug concentrations and/or timing of treatments were altered, as described in the Results. “Early pulse” treatment was from 24-48 hpf while “delayed” treatment was from 96-168 hpf (Fig. 4B). To obtain tissue for genotyping following treatments, larval heads were removed with a scalpel at the level of the pectoral fins. Larval tails with trunk skeletal muscle remained intact and were maintained in 10% formalin in PBS until imaging was performed.

Validation treatments at the University of Maine were carried out as described above except for the following modifications: Embryos were raised in a 28°C incubator with a 14 hr on, 10 hr off light cycle. At 24 hpf, embryos were manually dechorionated with forceps and sorted into 12-well plates at a density of 4-6 embryos/well. Vehicle control was 0.1% DMSO, while working concentration of SR-4370 remained at 1 µM. At 96 hpf, larvae were fixed in 4% paraformaldehyde (PFA) and stored at 4°C for 48 hours. Larvae were then rinsed in PBS-0.1% Tween 20 prior to imaging.

### 2.4. Imaging and scoring muscle lesions

Qualitative and quantitative scoring of *dmd* zebrafish larval muscle lesions was performed as previously described [17]. Birefringence was used to qualitatively sort and score larvae based on the binary presence (muscle lesions vs. no muscle lesions) and ordinal severity (unaffected/no lesions or mild, moderate, or severe lesions) of dystrophic muscle disease (Fig. 1C) [17], [35]. Quantitative measurements of birefringence were performed by mounting larvae in 2.5% methyl cellulose and adjusting camera and software settings (SZX16 stereomicroscope, DP72 camera, cellSens Dimension v4.1; Olympus Life Sciences, Bethlehem, PA) so that grayscale images had few to no saturated pixels across the entirety of the trunk, as previously described [17]. Average pixel intensity (*i.e.*, mean gray value) across the trunk muscle birefringence of individual larvae was calculated, and values were then normalized to the WT +DMSO control average values

### 2.5. Pool-specific risk ratio and composite scoring to obtain inferred rescue effect of individual compounds

Because of the number of treatments and animals needed, screening of the 185 drug pools of the epigenetic library was performed over multiple batches and zebrafish breedings. Therefore, frequencies of affected animals (using binary scoring) across chemical treatment pools were normalized relative to the batch-specific vehicle control average. These normalized results were then fitted to a logistic regression model (R package: GLM, v14) that assigned each drug pool a risk ratio reflective of the likelihood of affected animals (Table S3).

To assign individual compounds an inferred rescue score based on the effects of their corresponding row and column drug pools, we developed a composite score that incorporated both binary scoring (affected vs. unaffected) and ordinal scoring (lesion severity), using the following formula (Formula 1):

**Formula 1**

Average percent mild for an exposed group or control treatment with *n* replicates:

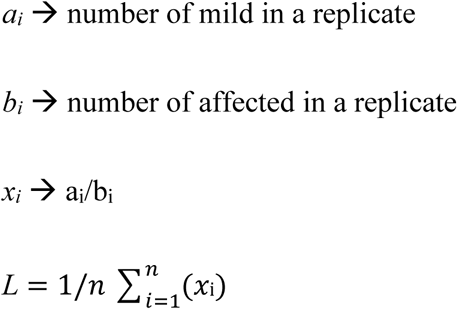

For a given exposed row or column group, difference from the corresponding batch DMSO control:

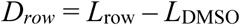

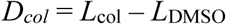

Inferred rescue from intersecting pools:

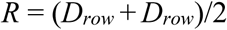

The composite score thus accounts for results from intersecting row and column drug pool treatments. Formula 1 values for individual compounds (composite scores) are provided in Table S4 and are plotted in Fig 2B.

**Fig. 2.**
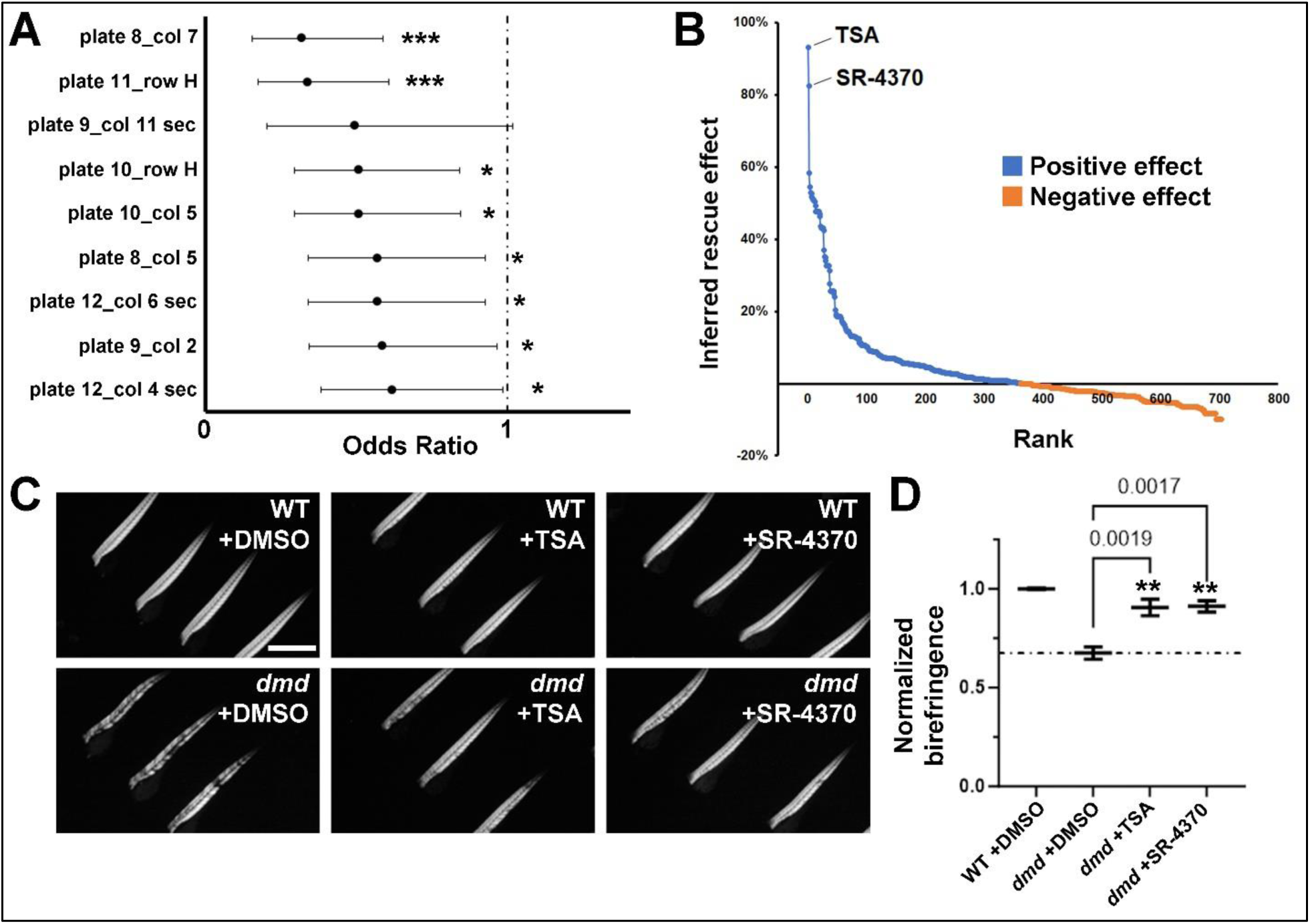
Composite scoring identifies the HDACi SR-4370 as a beneficial compound. **A.** Drug treatment pools with the lowest median odds of lesion development. Batch-corrected median odds ratios (OR) were calculated relative to DMSO control treatments. Median odds ratio is shown by a closed circle and whiskers represent the 95% confidence intervals. OR<1 indicates reduced chance of lesion development. Secondary pools (designated by the suffix “_sec”) refer to pools with suspected toxic compounds removed. Only one combination of the 8 pools with significantly reduced odds of lesion development intersected upon a compound (molibresib, [plate 10_row H], [plate 10_col 5]). *** P<0.0007. * P<0.05. **B.** Distribution of inferred rescue effects for individual compounds, as calculated using Formula 1. The two highest ranked compounds are HDACi trichostatin A (TSA) and SR-4370. **C.** Lateral views of trunk muscle birefringence in 96 hpf wild type (WT; *dmd+/+*) and *dmd* (*dmd−/−*) animals treated with DMSO, 1 µM TSA, or 1 µM SR-4370 from 24-96 hpf. Scale bar = 1mm. **D.** Normalized birefringence pixel intensities for DMSO, 1 µM TSA, and 1 µM SR-4370 treatments from 24-96 hpf. n=4 replicates for each treatment. Numbers of animals (per replicate and total ∑ across all 4 replicates) for each treatment and genotype combination are as follows: WT +DMSO (4-5, ∑=19), *dmd* +DMSO (4-9, ∑=28), *dmd* +TSA (2-6, ∑=17), *dmd* +SR-4370 (3-5, ∑=19). P values (**) are shown compared to *dmd* +DMSO control (dashed line).

### 2.6. Immunoblotting

Following HDACi exposure from 24-48 hpf, 25 larvae/replicate were de-yolked by rinsing in cold Ringer’s solution then lysed in 12.5 µL/larva of 1.5x LDS sample buffer (Cat. No. NP0007; Invitrogen, Waltham, MA). Lysates of 1 larva equivalent were then separated on reducing 12% NuPage Bis-Tris gels (NP0342BOX; Invitrogen), run in NuPAGE MES SDS buffer (Cat. No. NP0002; Invitrogen), and blocked in LI-COR TBS Intercept block (Cat. No. 927-660003; LI-COR Biosciences, Lincoln, NE). Blots were probed using the following antibodies: anti-acetyl-histone H3 (K9) (1:4000; Cat. No. 07-352; MilliporeSigma, Burlington, MA), and anti-acetyl-histone H4 (recognizes acetylated K5, K8, K12, and K16); 1:2000; Cat. No. 06-866; MilliporeSigma). Anti-alpha-actin (1:4000; Cat. No. 0869100; MP Biomedical, Irvine, CA) was included as a loading control. Infrared-labeled secondary antibodies (Anti-rabbit IgG DyLight 800, Cat. No. 611-145-002, used at 1:10,000, and Anti-mouse IgG DyLight 680, Cat. No. 610-144-002, used at 1:20,000; Rockland Immunochemicals, Philadelphia, PA) were visualized using an Odyssey infrared imager (LI-COR Biosciences) and quantified using Image Studio Lite (version 5.2). See Fig. S3 for uncropped images of original blots.

### 2.7. Statistical analyses

Unless otherwise noted, data was analyzed using a one-way ANOVA comparing each treatment group/condition to the *dmd* DMSO control group, with Dunnett’s correction for multiple comparisons, and a cutoff of p < 0.05. Unless otherwise noted, error bars represent standard error of the mean between replicates. Graphs and statistics were generated in GraphPad Prism (version 10.0.3).

## 3. Results

### 3.1. Drug-pool-based epigenetic library screening identifies beneficial drug pools for dmd zebrafish

In our previous work, we used a pilot screen of a commercial 94-chemical epigenetic small molecule library to identify a combination of two HDACi compounds, oxamflatin and salermide, that ameliorated *dmd* mutant zebrafish skeletal muscle degeneration [17]. Here, we sought to test whether we could identify additional novel epigenetic small molecules that improve zebrafish *dmd* muscle lesions. We selected the commercially available MedChemExpress Epigenetics Compound Library. The version of the library that we obtained contained 817 chemicals distributed over fourteen 96-well plates (Table S1). To efficiently screen this large library, we used an orthogonal drug pooling approach (Fig. 1A). We divided each library plate into plate-specific row- and column-based drug pool combinations, resulting in 185 unique pools across the library. Each compound was thus included in two separate pools: a row pool and a column pool (Fig. 1A).

Based on our previous work [17], we used a dose of 1 μM of each compound in a row or column pool for *dmd* zebrafish screening treatments. Embryos from *dmd+/−* crosses were treated from 24-96 hours post fertilization (hpf) (Fig. 1B). Each row- and column-based drug pool was tested in quadruplicate, with 25 animals per well for each replicate (Fig. 1B). After treatments, animals were fixed to preserve them for scoring muscle birefringence, whereby polarized light transmission appears reduced in the damaged muscle of diseased animals (Fig. 1B-1C). Our approach for assessing the dystrophic phenotype in this drug screen follows the approach that we and others used in previous zebrafish *dmd* chemical screens [17], [29], [36], [37]. In this approach, animals are scored as “affected”, showing a *dmd* mutant (*dmd−/−*) muscle lesion phenotype, or “unaffected”, showing normal birefringence (Fig. 1C).

During our screening, general embryo lethality (unrelated to *dmd*) was observed for 51 of the 185 unique treatment pools, thereby precluding birefringence-based phenotypic scoring of larvae at 96 hpf. Within the corresponding DMSO-vehicle control groups, only low levels of mortality occurred (Fig. S1). The intersections of the 51 “lethal” row and column pools identified 71 compounds (“excluded compounds” in Table S1). We thus hypothesized that we could circumvent the embryonic lethality by excluding these 71 compounds from their respective drug pools. Following the exclusion of the 71 suspected toxic compounds, re-screening of the 51 pools showed that only 5 pools remained toxic, thus allowing birefringence scoring of 46 of the 51 previously lethal pools. Overall, 180 unique pools, and about 86% (703/817) of compounds in the library, were evaluated in ≥2 (initial or secondary row and column) treatment pools (Fig. 1D; Table S1).

For the 180 scoreable treatment pools, we determined the average percentage of affected animals from each pool treatment and from each corresponding DMSO control treatment. By fitting the normalized binary (affected vs. unaffected) phenotypic scoring results to a logistic regression model, we determined that the odds of observing affected animals were significantly reduced in 8 of 180 pools (Fig. 2A; full results in Table S3). The pool with the second lowest odds of affected animals, [plate 11_row H], is noteworthy because it contains trichostatin A (TSA), a pan-HDAC inhibitor previously shown to improve mouse and zebrafish DMD models [16], [17], [19]. However, out of these top ranked pools, only one compound, molibresib, was identified through an intersecting pool ([plate 10_row H], [plate 10_col 5]; Fig. 2A). The inability to identify multiple positive row- and column-intersecting compounds from this logistic regression model suggests that requiring both row and column drug pool treatments to exhibit rescue is too stringent, as some drug pools may have inhibitory interactions. In addition, binary scoring of larvae as affected/unaffected may be insensitive to drug pool treatments that cause modest or incomplete rescue in *dmd* mutants.

### 3.2. Composite scoring identifies the HDACi SR-4370 as a beneficial compound

In order to take into account both row and column treatment effects, as well as treatments that cause a reduction in lesion severity without achieving complete rescue, we developed a composite scoring approach that allowed us to incorporate both ordinal scoring (unaffected, mild, moderate, or severe; see Fig. 1C) and binary scoring (percentage of affected animals) for both row and column treatments (see Formula 1 in Materials and methods). In control DMSO treatments, about 25% of larvae from *dmd+/−* crosses appear affected and exhibit muscle lesions predominantly scored as severe at 96 hpf (Fig. S1). Compounds in row and/or column pool treatments with <25% affected larvae and/or a greater proportion of mild vs. severe lesions were calculated as having higher composite scores (Table S4). We used these composite scores to generate a ranked plot of the inferred rescue effects of individual library compounds (Fig. 2B). TSA emerged as the compound with the greatest inferred rescue effect (93% average percent increase in larvae with mild lesions in [plate 11_row H] and [plate 11_col 6] treatments compared to DMSO controls) (Fig. 2B; Table S4), supporting the utility of using our formula to rank the rescue effects of individual compounds.

The composite scoring identifies the HDACi SR-4370 as the compound with the second highest inferred rescue effect (83% average percent increase in larvae with mild lesions compared to DMSO control) (Fig. 2B, Table S4). This score primarily reflected SR-4370’s inclusion in the pool [plate 8_col 7], which had the overall lowest odds of dystrophic lesion development (median OR=0.319; Fig. 2A; Table S3). We therefore tested SR-4370 for its ability to rescue *dmd* lesions as an individual compound. 1 μM treatments of SR-4370, from 24-96 hpf, lead to significantly improved *dmd* birefringence (Fig. 2C-D). We then tested additional individual compounds from the composite scoring, including the next top 10 compounds with the highest composite scores, as well as molibresib (see Fig. 2A) and other compounds with a range of composite score ranks (Fig. S2). Aside from TSA and SR-4370, all other individual compounds tested show no significant effects on *dmd* rescue (Fig. S2). The lack of effect of additional compounds correlates with the steep reduction in inferred rescue effect between SR-4370 (rank 2) and the next highest ranked compound (CC-90005, rank 3) (Fig. 2B and Table S4). Thus, through pool-based screening and our composite scoring approach, we identified SR-4370 as a novel HDACi compound that improves zebrafish *dmd* muscle lesions.

### 3.3. Additional validations corroborate the beneficial effects of SR-4370

Although the effects of SR-4370 and SR-4370-containing treatment pools were consistent across multiple experimental batches, all compounds were sourced from a single vendor, thereby representing a potential source of confirmation bias. We therefore validated the source-independent effects of SR-4370 by ordering drug from 3 different vendors (Table S2) and tested them using standard 1 μM treatments from 24-96 hpf. *dmd* muscle birefringence was significantly improved across all treatments of SR-4370 from multiple sources (Fig. 3A). Normalized birefringence ranged between about 82%-88%, indicative of consistent phenotypic improvement across groups (Fig. 3A).

**Fig. 3.**
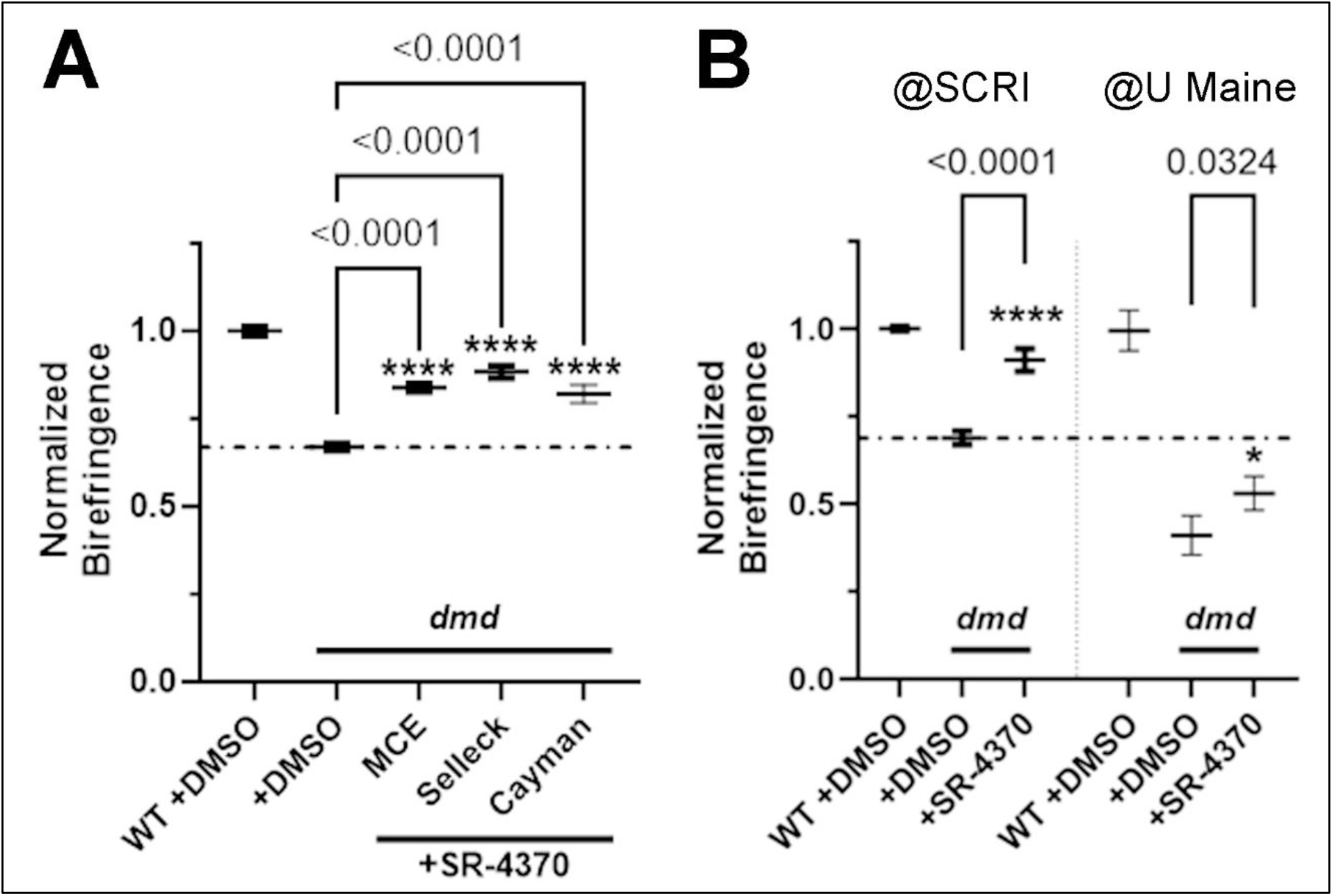
*dmd* rescue by SR-4370 is independent of chemical source and treatment site. **A.** Normalized birefringence pixel intensities for animals exposed to 1 µM SR-4370 from 24-96 hpf. Compounds were obtained from multiple sources (MCE, MedChemExpress; Selleck Chemicals, Houston, TX; Cayman Chemicals, Ann Arbor, MI). n=4 replicates for each treatment Numbers of animals (per replicate and total across all replicates) for each treatment and genotype combination are as follows: WT +DMSO (6-9, ∑=27), *dmd* +DMSO (4-7, ∑=24), *dmd* +MCE (4-9, ∑=26), *dmd* +Selleck (6-10, ∑=30), *dmd* +Cayman (6-7, ∑=25). P values (****) are shown compared to *dmd* +DMSO control (dashed line). **B.** Independent lab site comparison of normalized birefringence pixel intensities for animals exposed to 1 µM SR-4370 from 24-96 hpf. These independent tests, performed at Seattle Children’s Research Institute (SCRI) and at University of Maine (U Maine) and separated by vertical dotted line on the graph, were separately normalized to their corresponding WT +DMSO pixel intensities. Numbers of animals for each treatment and genotype combination are as follows: For SCRI: WT +DMSO (n=19), *dmd* +DMSO (n=24), *dmd* +SR-4370 (n=18); For UMaine: WT +DMSO (n=18), *dmd* +DMSO (n=15), *dmd* +SR-4370 (n=20). Error bars represent standard deviation. Significance was determined using a one-way ANOVA comparing each treatment group to their *dmd +*DMSO control group. P values are shown compared to the respective *dmd* +DMSO controls.

To independently validate the effects of SR-4370, we performed treatments at a different site, at the University of Maine, using standard 1 μM SR-4370 treatments from 24-96 hpf. Although normalized *dmd* birefringence was lower in animals at University of Maine compared to that at Seattle Children’s Research Institute, *dmd* muscle was significantly improved at both facilities (Fig. 3B). By demonstrating reproducibility, these independent tests are a strong validation of the benefits of SR-4370 in *dmd* mutant zebrafish.

### 3.4. HDACi treatments on dmd embryos are effective prior to muscle lesion formation

We next sought to determine the optimal SR-4370 treatment dose and timing necessary for *dmd* muscle lesion improvement. We tested doses of SR-4370 from 0.1-10 µM using the standard 24-96 hpf treatment conditions. We observe a dose response between 0.1-1 µM, with 500 nM being the minimum threshold for effective rescue of birefringence (Fig. 4A). 2.5 µM treatment does not significantly improve *dmd* muscle birefringence and causes substantial lethality, and embryonic lethality (of both WT and *dmd* embryos) occurs with 5 µM and 10 µM treatments (Fig. 4A). These results identify 1 µM as the optimal dose for 24-96 hpf treatments.

**Fig. 4.**
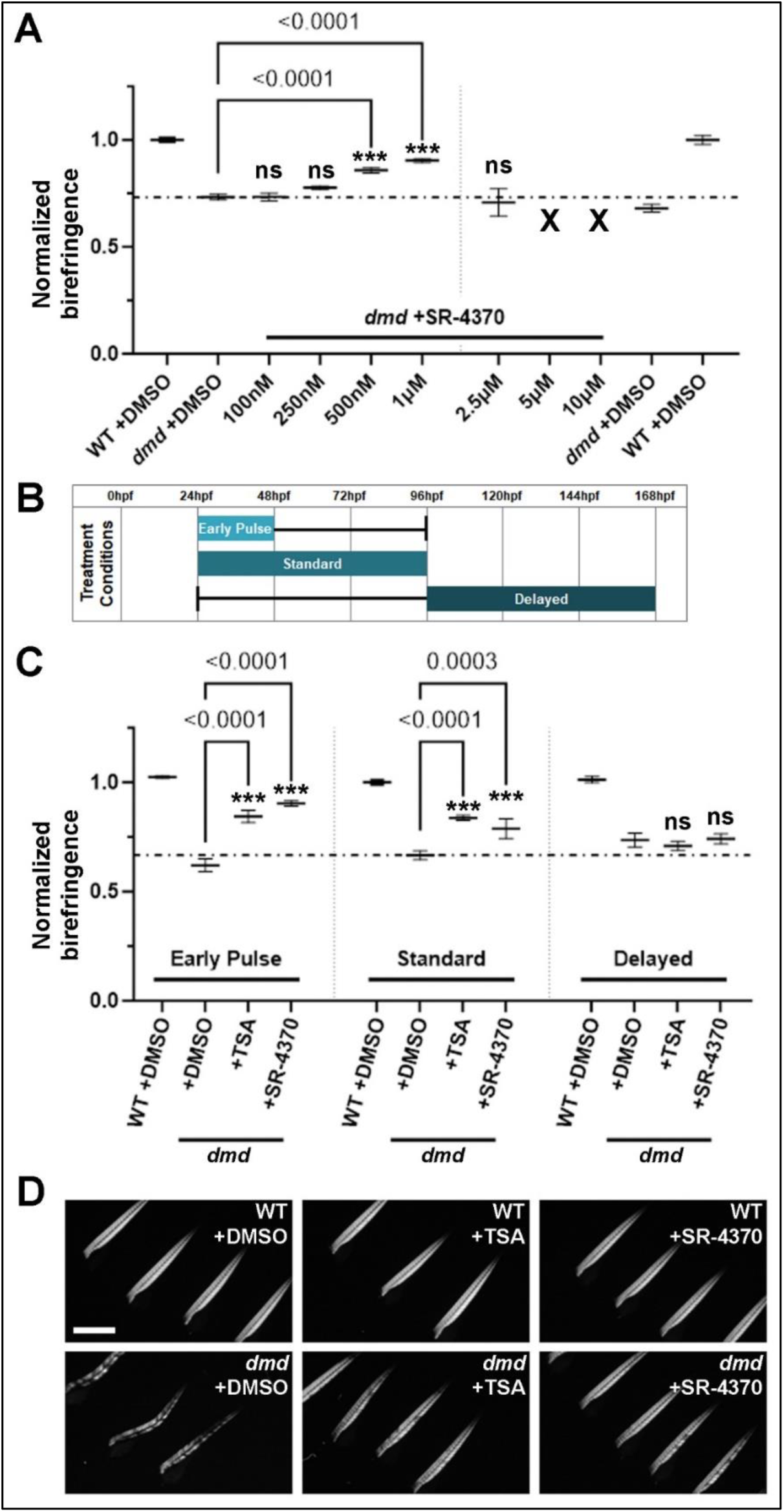
*dmd* rescue by SR-4370 follows a dose-response pattern and occurs during an early treatment window. **A.** Normalized birefringence pixel intensities for animals exposed to 100 nM-10 µM SR-4370 from 24-96 hpf. For 100 nM-1 µM treatments, n=4 replicates for each treatment. Mortality in the 2.5 µM treatments reduced counts to n=3 replicates. Complete lethality was seen in the 5 µM and 10 µM groups (indicated by X). Numbers of animals (per replicate and total across all replicates) for each treatment and genotype combination are as follows: WT +DMSO (5-6, ∑=22), *dmd* +DMSO (5-8, ∑=26), *dmd* +100nM (3-6, ∑=20), *dmd* +250nM (6-10, ∑=31), *dmd* +500nM (5-7, ∑=24), *dmd* +1µM (4-8, ∑=21), *dmd* +2.5µM (1-4, ∑=8). The 100 nM-1 µM treatments and the 2.5 µM-10 µM treatments were performed separately (shown with vertical dotted line) and thus their corresponding DMSO controls are shown. P values (***) are shown. ns = not significant. Horizontal dashed line indicates *dmd* +DMSO level (for 100 nM-1 µM treatments). **B.** Timeline of alternate small molecule treatment conditions. For early pulse, animals were exposed from 24-48 hpf and then reared in plain embryo media (see Methods) until 96 hpf. Media and compounds were replaced daily for standard and delayed treatments. At end points, animals were fixed and then scored using birefringence. **C.** Normalized birefringence pixel intensities for animals exposed to 200 nM TSA or 1 µM SR-4370 from 24-48 hpf, 24-96 hpf, or 96-168 hpf. n=4 replicates for each treatment. Numbers of animals (per replicate and total across all 4 replicates) for each treatment and genotype combination are as follows: early pulse WT +DMSO (4-7, ∑=24), *dmd* +DMSO (3-6, ∑=16), *dmd* +TSA (4-10, ∑=25), *dmd* +SR-4370 (5-11, ∑=27); standard WT +DMSO (1-7, ∑=16), *dmd* +DMSO (1-9, ∑=22), *dmd* +TSA (4-12, ∑=30), *dmd* +SR-4370 (3-10, ∑=21); delayed WT +DMSO (5-9, ∑=26), *dmd* +DMSO (3-7, ∑=20), *dmd* +TSA (2-7, ∑=19), *dmd* +SR-4370 (6-7, ∑=26). P values (***) are shown. ns = not significant. Horizontal dashed line indicates *dmd* +DMSO level (for 24-96 hpf treatments). **D.** Lateral views of trunk muscle birefringence in 96 hpf WT (*dmd+/+*) and *dmd* (*dmd−/−*) animals treated with DMSO, 200 nM TSA, or 1 µM SR-4370 from 24-48 hpf. Scale bar =1mm.

Muscle lesions in *dmd* zebrafish are initially visible at 48 hpf but become progressively more conspicuous through 96 hpf [25]. We evaluated whether shortened administration of beneficial compounds prior to lesion formation could affect muscle morphology at later stages (Fig. 4B). Using birefringence as a readout, we observe that an abbreviated 24-48 hpf administration of either SR-4370 or TSA significantly improves the dystrophic phenotype at 96 hpf (Fig. 4C-D). We next tested whether our standard 3-day treatment length could be applied after lesion formation to improve *dmd* birefringence. We find that treatments from 96-168 hpf do not provide any detectable improvement of *dmd* birefringence (Fig. 4C). These results suggest that HDACi compounds are effective during a critical early window of zebrafish development.

### 3.5. Drug pool screening identifies complex drug combinations beneficial for dmd zebrafish

Because our previous studies identified the HDACi oxamflatin as beneficial for *dmd* zebrafish, we re-examined oxamflatin-containing drug pools from the current library screen to determine whether there was evidence of phenotypic improvement. Oxamflatin is located at library position [plate 7_F10] (Table S1) and only achieved a composite rescue effect of 13% (Table S4). Pool [plate 7_col 10] had slightly decreased odds of lesion development (median OR=0.725), while the corresponding row pool containing oxamflatin, [plate 7_row F], demonstrated increased odds of lesion development (median OR=1.290) (Table S3). We assessed quantitative birefringence from these two pool treatments and found that [plate 7_row F] did significantly improve *dmd* birefringence (Fig. 5A). However, individual compound testing revealed that none of the 10 compounds in pool [plate 7_row F] significantly improved *dmd* muscle birefringence (Fig. 5B). These results therefore suggest that a combination of drugs within pool [plate 7_row F] is contributing to the rescue effects of this drug pool.

**Fig. 5.**
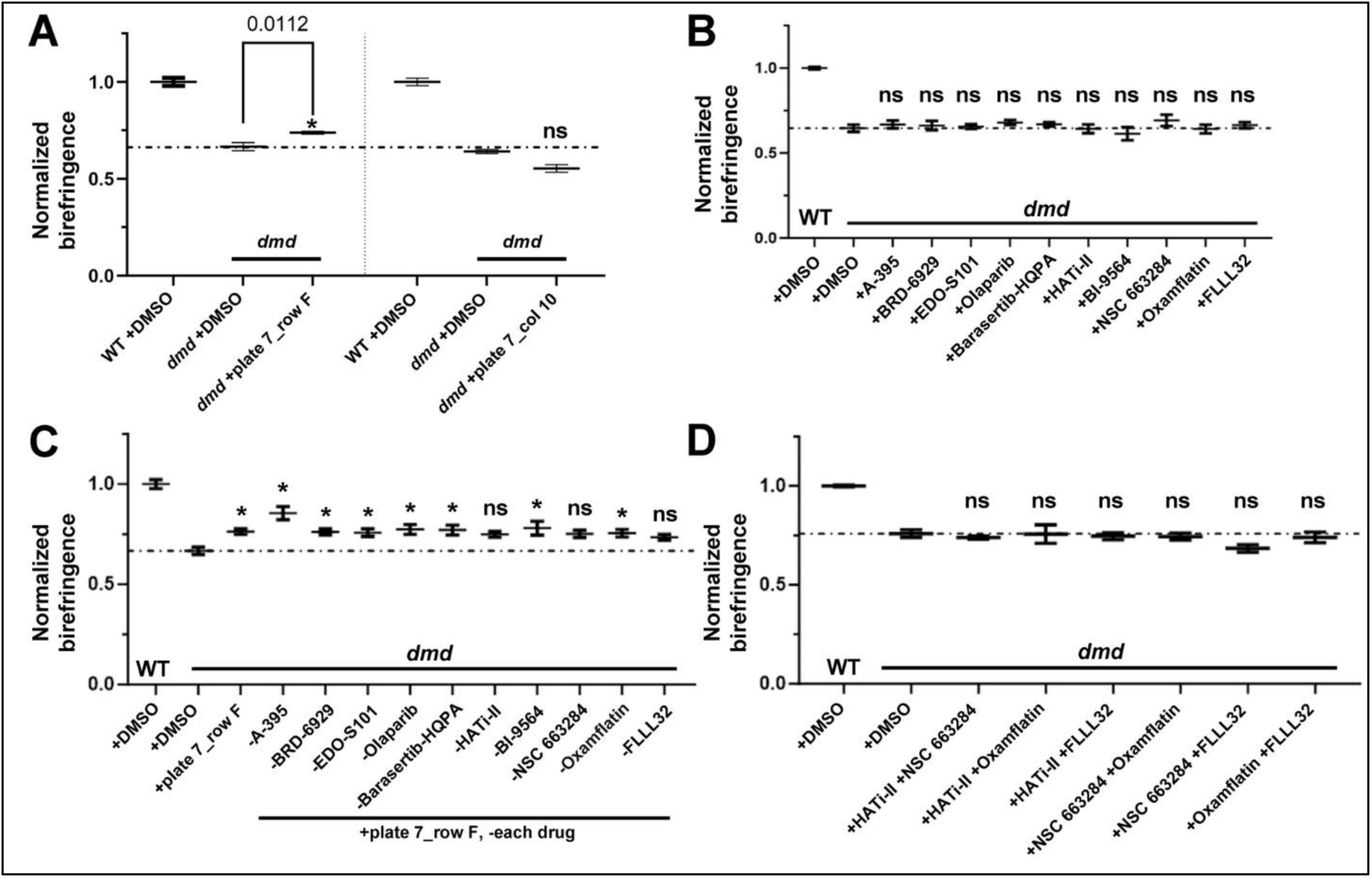
Drug combinations containing the HDACi oxamflatin are beneficial for *dmd* zebrafish. **A.** Normalized birefringence pixel intensities for 24-96 hpf treatments with DMSO, pool [plate 7_row F], and pool [plate 7_column C]. n=4 replicates for each treatment. Numbers of animals (per replicate and across all 4 replicates) for each treatment and genotype combination are as follows: WT +DMSO (5-10, ∑=31), *dmd* +DMSO (4-10, 28), *dmd* +plate7_row F (5-9, ∑=30); WT +DMSO (3-9, ∑=24), *dmd* +DMSO (3-9, ∑=25), *dmd* +plate7_col 10 (2-6, ∑=16). **B.** Normalized birefringence pixel intensities for 24-96 hpf treatments with DMSO and the 10 individual compounds in pool [plate 7_row F]. n=4 replicates for each treatment. Numbers of animals (per replicate and across all 4 replicates) for each treatment and genotype combination are as follows: WT +DMSO (3-8, ∑=22), *dmd* +DMSO (2-6, ∑=18), *dmd* +A-395 (4-6, ∑=19), *dmd* +BRD-6929 (5-11, ∑=35), *dmd +*EDO-S101 (3-9, ∑=25), *dmd* +Olaparib (3-8, ∑=24), *dmd* +Barasertib-HQPA (3-10, ∑=28), *dmd* +HATi-II (5-9, ∑=26), *dmd* +BI-9564 (3-8, ∑=21), *dmd* +NSC663284 (4-10, ∑=25), *dmd* +Oxamflatin (6-11, ∑=35), *dmd* +FLLL32 (4-8, ∑=23). **C.** Normalized birefringence pixel intensities for 24-96 hpf treatments with DMSO, pool [plate 7_row F], and [plate 7_row F] pools with each individual compound removed. n=4 replicates for each treatment. Numbers of animals (per replicate and across all 4 replicates) for each treatment and genotype combination are as follows: WT +DMSO (3-13, ∑=28), *dmd* +DMSO (2-6, ∑=19), *dmd* +plate 7_row F (2-6, ∑=14), *dmd -*A-395 (5-9, ∑=27), *dmd* -BRD-6929 (6-10, ∑=30), *dmd* -EDO-S101 (6-11, ∑=31), *dmd* -Olaparib (6-11, ∑=30), *dmd* -Barasertib-HQPA (2-6, ∑=16), *dmd* -HATi-II (5-11, ∑=31), *dmd* -BI-9564 (3-6, ∑=16), *dmd* -NSC-663284 (6-9, ∑=30), *dmd* - Oxamflatin (5-8, ∑=27), *dmd*-FLLL32 (3-10, ∑=26). **D.** Normalized birefringence pixel intensities for 24-96 hpf treatments with DMSO and pairwise combinations of HATi-II, NSC 663284, oxamflatin, and FLLL32. n=4 replicates for each treatment. Numbers of animals (per replicate and across all 4 replicates) for each treatment and genotype combination are as follows: WT +DMSO (3-7, ∑=21), *dmd* +DMSO (3-7, ∑=21), *dmd* +plate 7_row F (3-5, ∑=17), *dmd +*HATi-II+NSC663284 (3-10, ∑=29), *dmd +*HATi-II+Oxamflatin (3-9, ∑=23), *dmd +*HATi-II+FLLL32 (6-7, ∑=25), *dmd +*NSC663284+Oxamflatin (4-8, ∑=23), *dmd +*NSC663284+FLLL32 (2-10, ∑=26), *dmd +*Oxamflatin+FLLL32 (2-6, ∑=17). For A.-D., horizontal dashed lines indicate *dmd* +DMSO levels. P values are shown: *p<0.05, **p<0.01, ***p<0.001, ****p<0.0001. ns = not significant.

We next sought to narrow down which compounds from [plate 7_row F] were necessary for improving *dmd* birefringence by systematically removing individual compounds from the drug pool. We thus tested a series of drug pools, containing 9 compounds each, and compared their effects to the full 10-compound pool from [plate 7_row F]. Three compounds, HATi-II, NSC 663284, and FLLL32, were implicated as necessary for the rescue effects of [plate 7_row F] since drug pools with these compounds removed did not significantly improve *dmd* muscle birefringence (Fig. 5C). However, pairwise combinations of these compounds, including combinations with oxamflatin, exhibited no rescue effects (Fig. 5D), indicating that a combination of greater than two compounds from [plate 7_row F] was likely necessary to improve the muscle phenotype in *dmd* mutants. Thus, even though in our previous studies we were able to identify a beneficial two-drug combination that includes oxamflatin, these results suggest multi-epigenetic compound combinations also have the potential to provide modest rescue of *dmd* birefringence.

### 3.6. Birefringence rescue by HDACi correlates with increased histone acetylation

We next sought to mechanistically validate and compare the effects of different HDACi treatments in *dmd* zebrafish. We first examined the effective doses of the HDACi TSA, oxamflatin, and givinostat for improving *dmd* muscle birefringence, using 24-48 hpf treatments (Fig. 6A-B). Consistent with our previous studies [16], [17], 200 nM TSA robustly improved *dmd* birefringence, whereas 1 µM oxamflatin did not induce any rescue (Fig. 6A-B). Both 10 µM and 20 µM oxamflatin treatments increased muscle birefringence (Fig. 6A-B). However, no birefringence improvement was seen with 20 nM TSA, or with any tested dose of givinostat (Fig. 6A). 200 µM givinostat treatments resulted in precipitation and the formation of a crystalline film in each well, suggesting that we exceeded the highest possible treatment dose for givinostat. These results reveal variation among HDACi treatments and effective doses in *dmd* zebrafish.

**Fig. 6.**
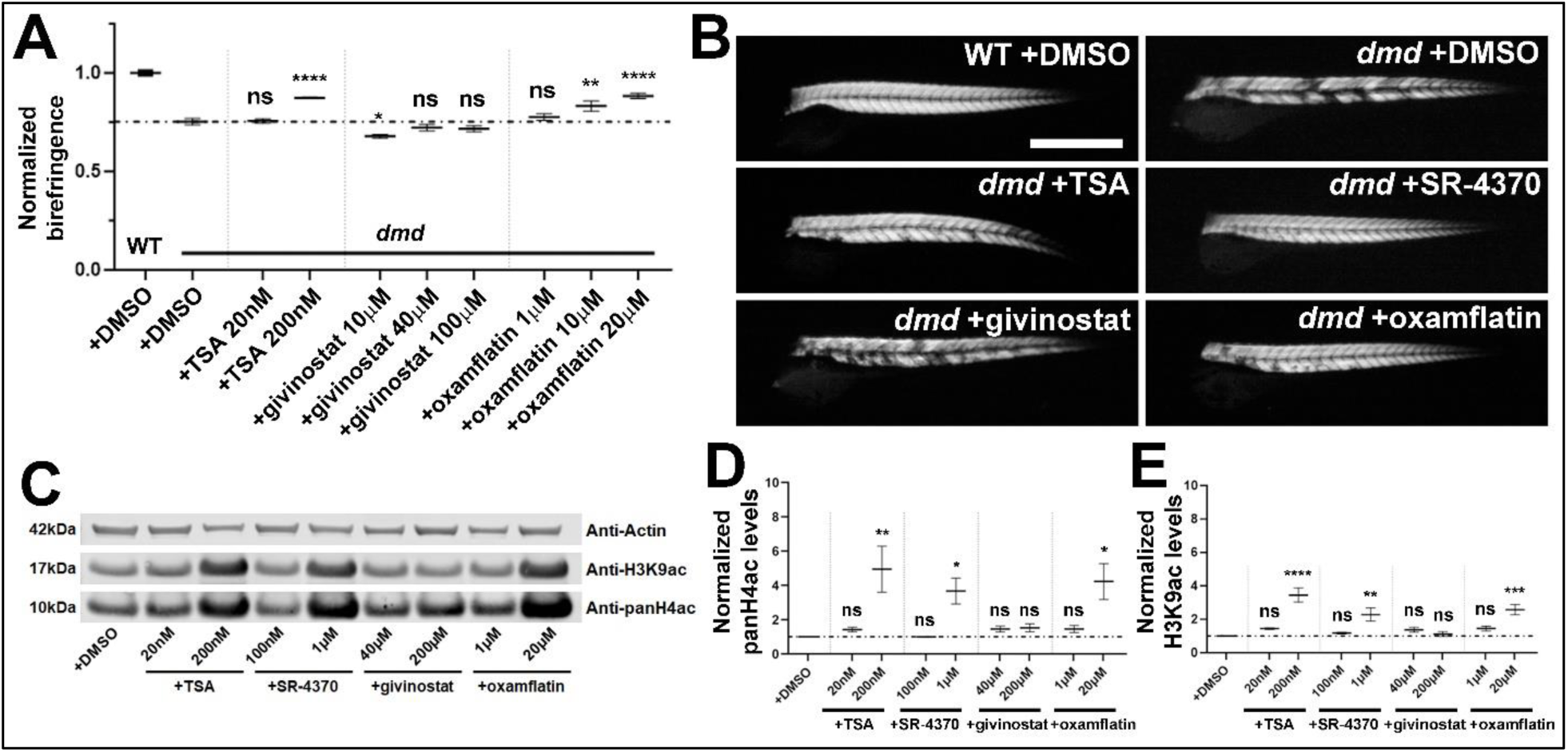
HDACi rescue correlates with increased histone acetylation. **A.** Normalized birefringence pixel intensities for animals exposed to doses of TSA, givinostat, or oxamflatin from 24-96 hpf. n=4 replicates for each treatment. Numbers of animals (per replicate and across all 4 replicates) for each treatment and genotype combination are as follows: WT +DMSO (4-7, ∑=24), *dmd* +DMSO (2-6, ∑=17), *dmd* +TSA 20nM (5-9, ∑=28), *dmd* +TSA 200nM (4-6, ∑=20), *dmd* +Givinostat 10µM (3-8, ∑=24), *dmd* +Givinostat 40µM (5-10, ∑=29), *dmd* +Givinostat 200µM (4-8, ∑=23), *dmd* +Oxamflatin 1µM (4-9, ∑=26), *dmd* +Givinostat 10µM (5-9, ∑=27), *dmd* +Givinostat 20µM (3-9, ∑=20). P values are shown: (*p<0.05, **p<0.01, ***p<0.001, ****p<0.0001). ns = not significant. **B.** Lateral views of trunk muscle birefringence in 96 hpf WT (*dmd+/+*) and *dmd* (*dmd−/−*) animals treated from 24-96 hpf with DMSO, 200 nM TSA, 1 µM SR-4370, 200 µM givinostat, or 20 µM oxamflatin. Scale bar =1mm. **C.** Western blots of 48 hpf larval lysates from animals treated with different HDACi doses, as in A. **D.-E.** Quantification of western blots (n=3). **D.** Pan-H4ac levels in non-genotyped 48 hpf larvae from *dmd+/−* clutches. **E.** H3K9ac levels in non-genotyped 48 hpf larvae from *dmd+/−* clutches. Pixel intensities were normalized to the anti-actin band in DMSO-treated animals. P values are shown: (*p<0.05, **p<0.01, ***p<0.001, ****p<0.0001). ns = not significant.

We next examined whether the effective and ineffective doses of these different HDACi led to altered histone acetylation levels. As pan-HDACi, TSA, SR-4370, givinostat, and oxamflatin are expected to induce increased histone H4 and histone H3 acetylation [17], [19], [23], [38], [39], [40], [41]. Western blot analysis revealed that both pan-H4ac and H3K9ac were significantly elevated in larvae exposed to: 200nM TSA, 1 µM SR-4370, and 20 µM oxamflatin (Fig. 6C-E). These changes correlated with HDACi doses that improved birefringence. Furthermore, we did not observe increased histone acetylation following givinostat treatments (40 and 200 µM) or ineffective doses of TSA, SR-4370, and oxamflatin (Fig. 6). Therefore, we conclude that HDACi treatments that improve the muscle phenotype of *dmd* zebrafish are strongly associated with increased histone acetylation.

## 4. Discussion

In this investigation, we used a drug-pooling approach to screen over 800 epigenetic small molecules for their ability to alleviate muscle lesions in the zebrafish *dmd* model. We identified a novel HDAC inhibitor (HDACi), SR-4370, that significantly improved muscle structure in *dmd* zebrafish larvae. After conducting dose-response experiments, we found that 500 nM and 1 µM doses of SR-4370 significantly ameliorated muscle lesion severity. Treatments with the HDACi SR-4370 and TSA significantly rescued the *dmd* phenotype when embryos were exposed from 24 to 48 hpf, prior to *dmd* muscle lesion appearance. However, a later exposure window of 96 hpf to 168 hpf did not result in an improved muscle phenotype. Furthermore, the ability of the HDACi TSA, SR-4370, and oxamflatin to improve *dmd* zebrafish muscle correlated with the ability of these HDACi to cause increased levels of histone acetylation in zebrafish larvae. Our findings support the use of zebrafish as an efficient screening tool for identifying small molecules with *in vivo* alleviatory effects in damaged muscle. Beyond novel identification of SR-4370 as a beneficial compound, our findings also provide further support for HDACi as a promising class of compounds for treating DMD.

Zebrafish are a well-established model for drug screening and drug discovery for human disease and, in particular, muscular dystrophy [30], [42], [43], [44]. The orthogonal drug pooling method that we employed here is similar to the approach used in a previous screen for antiangiogenic compounds in zebrafish larvae [45]. Pooled chemical screens can save time and resources as well as significantly reduce the number of animals used relative to single compound efforts. One limitation of the pooled-compound approach is that overly toxic individual chemicals can cause drug pools to be unfeasible to screen. We addressed this issue by orthogonally identifying likely toxic candidate compounds, removing these compounds, and rescreening the drug pools. Another limitation of the pooled-compound approach is that pooled compounds may inhibit or mask each other’s activity without apparent toxicity. We addressed this issue by generating a composite scoring approach, which allowed us to identify SR-4370, based largely on its activity in its column pool. It is possible that we failed to identify other beneficial compounds in the library because their activities were inhibited in both their row and column pools. However, aside from the positive effects shown here for the HDACi TSA, SR-4370, and oxamflatin, we did test 37 additional compounds individually during our study (see Fig. S2 and Fig. 5) and found no clear *dmd* rescue effects from these compounds.

One of our key findings is that HDACi treatments are effective at improving zebrafish *dmd* birefringence when they are initiated at an embryonic stage prior to overt *dmd* muscle lesion formation, but treatments are not effective when initiated after significant lesion formation. These results are consistent with previous studies showing that young (1.5 or 3 month old) *mdx* mice treated with HDACi show improved muscle morphology and function, including increased myofiber size and integrity, while old (12 month old) *mdx* mice are resistant to the beneficial effects of HDACi [19], [20], [46], [47]. The age-dependent effects of HDACi in *mdx* mice are likely mediated by the age- and disease-dependent plasticity of fibro-adipogenic progenitors (FAPs), which are multipotent muscle mesenchymal cells [23], [41], [46], [47]. In early-stage *mdx* mice, HDACi can induce FAPs toward a pro-myogenic and proliferative phenotype, whereas late-stage FAPs are resistant to these HDACi effects [23], [41], [46], [47], [48]. The effectiveness of HDACi during zebrafish embryogenesis may also be related to the stage-dependent ability of HDACi to promote myogenesis and muscle differentiation [39]. We do not yet know what cell type(s) HDACi are working through to improve *dmd* zebrafish.

A second key finding here is that the ability of HDACi treatments to improve *dmd* zebrafish muscle structure correlates with the ability of HDACi to cause increased histone acetylation in zebrafish larvae. We show that in zebrafish treated with the HDACi TSA, SR-4370, and oxamflatin, levels of H3K9 and pan-H4 acetylation were increased relative to control animals. We also previously showed that a combination treatment of the HDACi salermide and oxamflatin caused an increase in pan-H4 acetylation and H4K16 acetylation [17]. HDACi administration similarly causes histone hyperacetylation in *mdx* mice [19], [41], [49].

While our findings add to the growing evidence that HDACi are beneficial for DMD, we were not able to observe any improvements in muscle structure in *dmd* zebrafish treated with the HDACi givinostat, which has been shown to improve muscle structure in DMD boys [21]. This may be due to such factors as the solubility of givinostat in the zebrafish media or the ability of the embryo to uptake givinostat from the media. The fact that we were unable to detect increased levels of histone H3 or histone H4 acetylation in givinostat-treated embryos suggests that the drug was not reaching pharmacologically active concentrations in the animals. A previous study showed that givinostat is able to improve muscle structure in a morpholino antisense knock-down model of zebrafish *dmd* [18]. It is possible that differences in the embryo media or in the zebrafish model used (morpholino vs *sapje* mutant) could contribute to the disparate results.

The efficacy of SR-4370 may be due to the classes of HDACs it inhibits and its fluorinated chemical structure. SR-4370 is reported to inhibit HDAC1, 2, 3, 6, and 8 [50]. HDAC2 down-regulation leads to improved muscle structure and function in *mdx* mice [49]. Inhibiting HDAC6 in *mdx* mice results in improved muscle phenotype through Smad3 acetylation and downregulation of transforming growth factor beta signaling [51]. HDAC8 inhibition in dystrophic zebrafish results in α-tubulin acetylation and improved cytoskeleton organization [18]. Fluorination of SR-4370 may aid its ability to efficiently inhibit HDACs by increased binding affinity. For example, a comparison of inhibitory activity of the HDACi vorinostat to that of several synthesized 1,2-difluoro versions showed that the fluorinated compounds had increased potency against HDAC1 and HDAC6 compared to the parent compound [52]. Fluorinated analogues of the class I HDAC inhibitor largazole had stronger inhibition towards class I HDACs than unmodified largazole [53].

One of the limitations of our study, in addition to the drug pooling issues discussed above, is that we have not assessed the pharmacokinetics of these HDACi compounds in zebrafish tissues. With respect to zebrafish drug screening, most negative results are hard to interpret because it is usually not known if the compounds are being absorbed by the animals. We address this limitation, in part, by assaying a biochemical readout of the compounds being tested, in this case histone acetylation levels in zebrafish larvae treated with HDACi. In cases where we are able to observe positive benefits of HDACi on *dmd* muscle structure, we also observe increased histone acetylation levels. As discussed above, the lack of increased histone acetylation in givinostat-treated embryos may indicate that its failure to improve muscle structure is due to it not reaching active concentrations in the embryo. Future studies could further address the pharmacokinetics and pharmacodynamics of HDACi in *dmd* zebrafish.

Our study identifies SR-4370 as a potential therapeutic HDACi that merits further investigation in other models of DMD. Thus far, there has been only one previous publication on SR-4370 [50]. This study used a screen of epigenetic compounds to discover that SR-4370 can reactivate latent HIV-1 reservoirs, thus identifying SR-4370 as a novel potential lead compound for HIV therapy[50]. Future studies will be needed to further assess the promise of SR-4370 as a therapeutic for DMD.

## Acknowledgements

We would like to thank the SCRI Office of Animal Care for caring for the zebrafish.

## Author roles

Conceptualization: LM; Data curation: KL, EH, GHFIII, AI, LM; Formal analysis: KL, EH, GHFIII, AI, AGP, LM; Funding acquisition: LM; Investigation: KL, EH, GHFIII, AI, AP, AM, CH, LM; Methodology: KL, EH, GHFIII; Project administration: KL, LM; Resources: LM; Supervision: CH, LM; Validation: KL, EH, GHFIII, AI, AP, AM; Visualization: KL, EH, GHFIII, LM; Roles/Writing - original draft: KL, EH, LM; and Writing - review & editing: KL, EH, GHFIII, AI, AGP, CH, LM.

## Figures and Figure legends

**Fig. S1.**
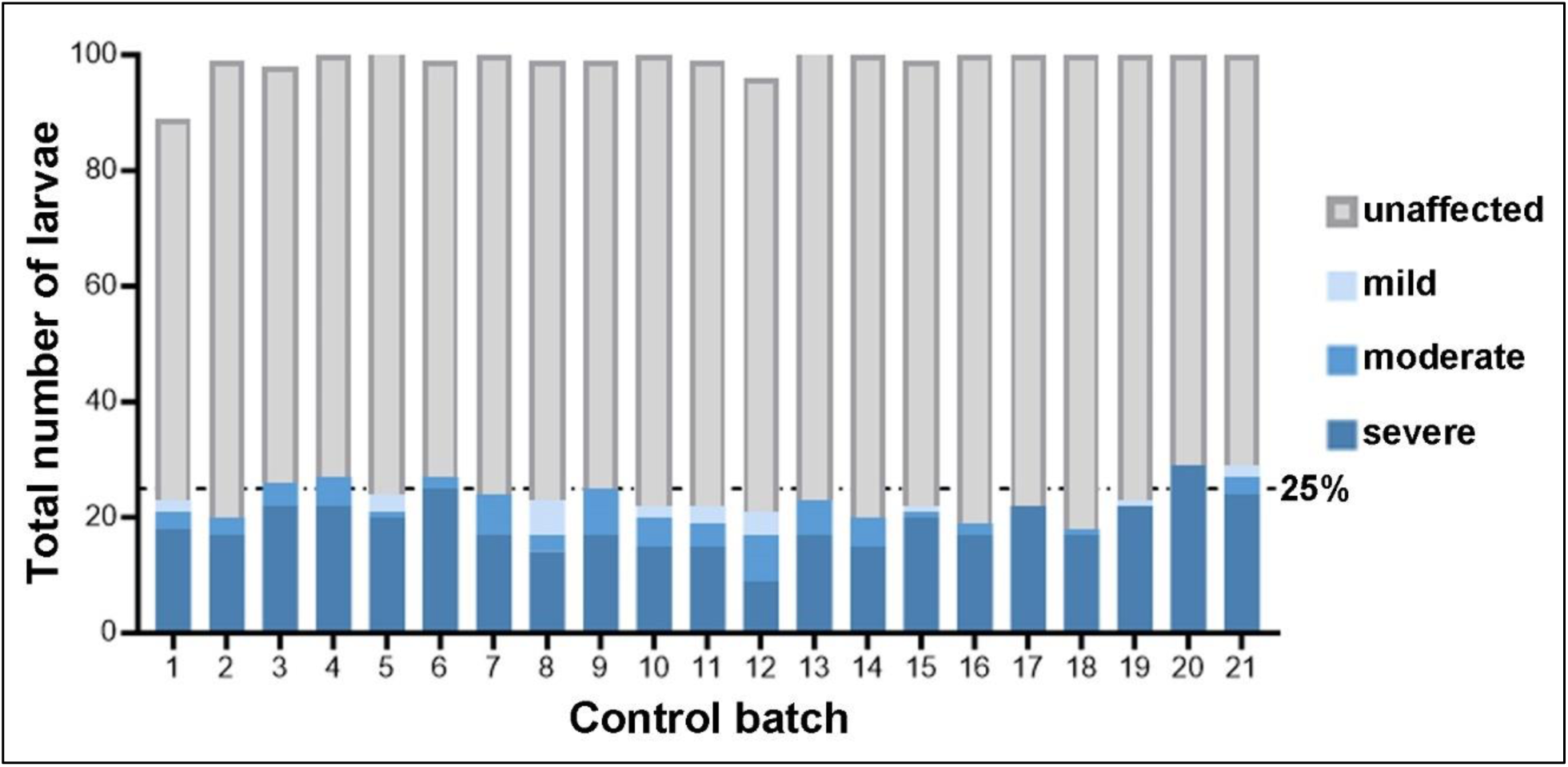
Larval lethality and proportion of affected animals is consistent across DMSO control groups. Screening of pooled compounds occurred over 21 different batches, each with a DMSO control (Control batch, x axis). Each control batch consisted of 4 wells of 25 animals each, with occasional loss of an animal (y axis). Stacked bars represent total qualitative ordinal scoring (unaffected, and mild, moderate, and severe affected) across 4 replicates. All DMSO control groups had ≥88% survival (average survival 98.9%±2.5 standard deviation [SD]) and followed an approximate Mendelian ratio of 25% (dashed line on y axis) affected individuals (average percent affected 23.5%±3.08 SD), as expected in the offspring from heterozygous *dmd+/−* crosses. Chi-squared test of observed distribution with the 25% expected Mendelian ratio showed no significant difference (p>0.9999) across all 21 batches. Larvae with severe lesions represented the majority (80.3%±14.2 SD) of affected larvae and also showed no significant difference (p>0.9999) across all 21 batches.

**Fig. S2.**
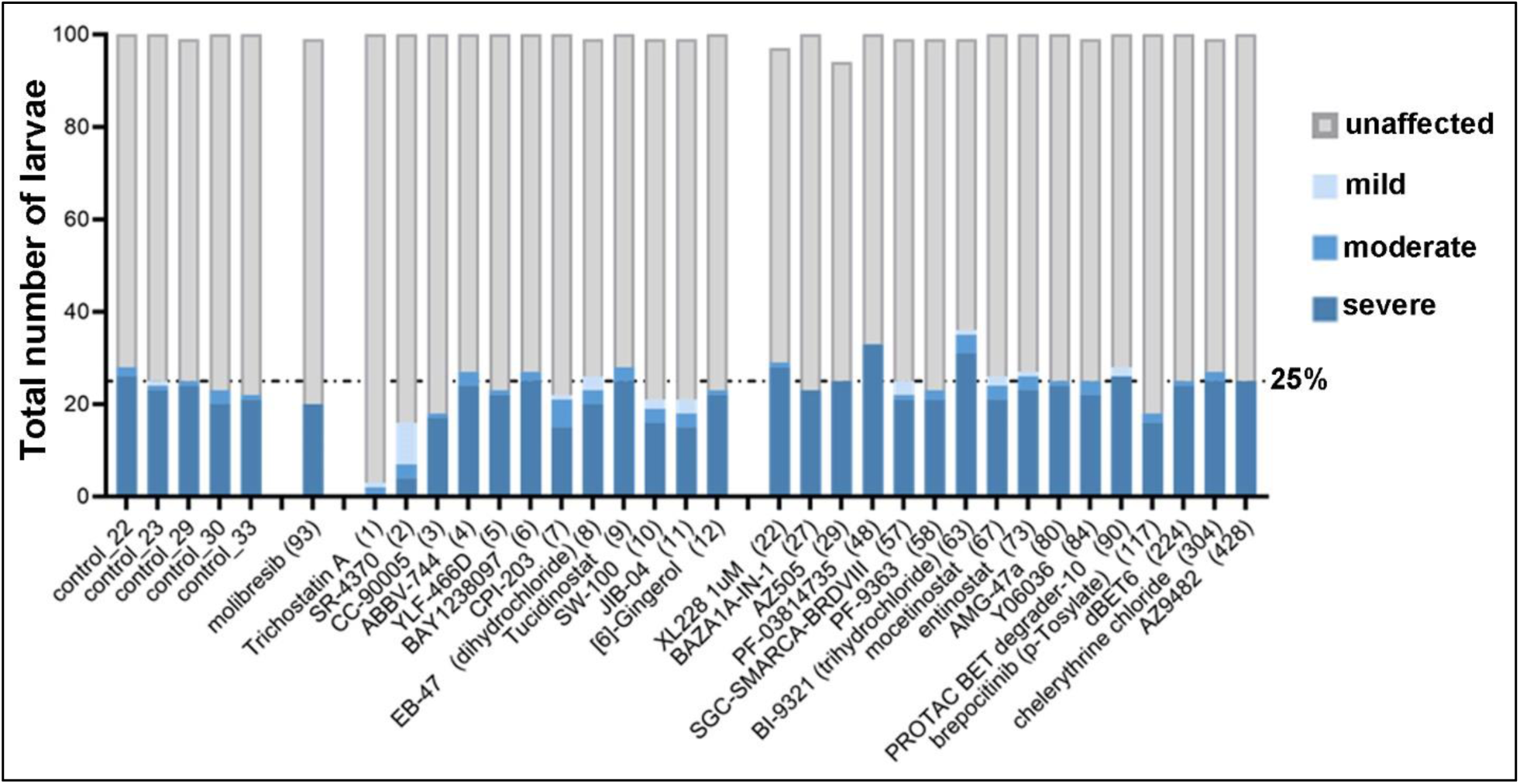
Validation tests of compounds based on composite scoring. Each control (DMSO) and treatment batch consisted of 4 wells of 25 animals each, with occasional loss of an animal (represented on y axis). Control treatments shown (with batch number) represent the control treatments performed for the batches encompassing the individual compounds tested. Composite scoring ranks for each compound are shown in parentheses. Stacked bars represent total qualitative ordinal scoring (unaffected, and mild, moderate, and severe affected) across 4 replicates. All control and treatment groups had ≥99% survival (average survival 98.6%±0.55 SD for controls and 99.4%±1.24 SD for treatments) and followed an approximate Mendelian ratio of 25% (dashed line on y axis) affected individuals (average percent affected 24.3%±2.90 SD for controls and 24.1%±5.90 SD for treatments). Larvae with severe lesions represented the majority (93.6%±4.85 SD) of affected larvae across all batches. TSA and SR-4370 showed lower percent of severe affected animals (0% and 25%, respectively) relative to controls and treatment group averages (93.6%±4.85 SD and 84.6%±21.99 SD).

**Fig. S3.**
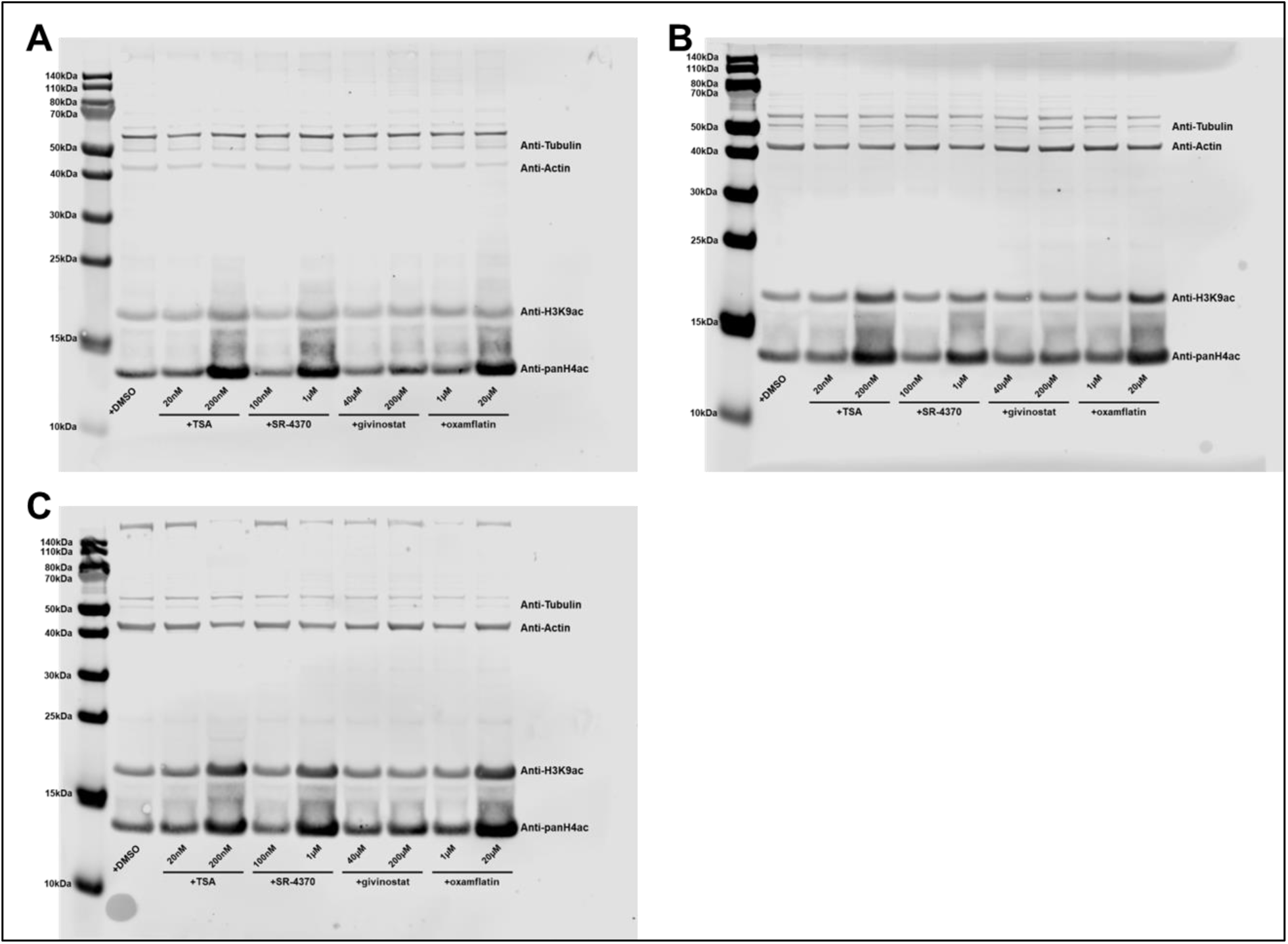
**A.-C.** Uncropped images of original Western blots used to create Fig. 6C-E. Each blot represents separate treatment replicates of the specified compounds and concentrations. Bands shown in Fig. 6C are taken from blot C.

## References

[1] E. P. Hoffman, R. H. Brown, and L. M. Kunkel, “Dystrophin: The protein product of the duchenne muscular dystrophy locus,” Cell, vol. 51, no. 6, pp. 919–928, Dec. 1987, doi: 10.1016/0092-8674(87)90579-4.

[2] A. H. M. Burghes, C. Logan, X. Hu, B. Belfall, R. G. Worton, and P. N. Ray, “A cDNA clone from the Duchenne/Becker muscular dystrophy gene,” Nature, vol. 328, no. 6129, pp. 434–437, Jul. 1987, doi: 10.1038/328434a0.

[3] D. Duan, N. Goemans, S. Takeda, E. Mercuri, and A. Aartsma-Rus, “Duchenne muscular dystrophy,” Nat Rev Dis Primers, vol. 7, no. 1, p. 13, Feb. 2021, doi: 10.1038/s41572-021-00248-3.

[4] A. M. Elasbali, W. A. Al-Soud, S. Anwar, H. H. Alhassan, M. Adnan, and Md. I. Hassan, “A review on mechanistic insights into structure and function of dystrophin protein in pathophysiology and therapeutic targeting of Duchenne muscular dystrophy,” International Journal of Biological Macromolecules, vol. 264, p. 130544, Apr. 2024, doi: 10.1016/j.ijbiomac.2024.130544.

[5] D. G. Allen, N. P. Whitehead, and S. C. Froehner, “Absence of Dystrophin Disrupts Skeletal Muscle Signaling: Roles of Ca2+, Reactive Oxygen Species, and Nitric Oxide in the Development of Muscular Dystrophy,” Physiological Reviews, vol. 96, no. 1, pp. 253–305, Jan. 2016, doi: 10.1152/physrev.00007.2015.

[6] P. Dowling, D. Swandulla, and K. Ohlendieck, “Cellular pathogenesis of Duchenne muscular dystrophy: progressive myofibre degeneration, chronic inflammation, reactive myofibrosis and satellite cell dysfunction,” Eur J Transl Myol, Oct. 2023, doi: 10.4081/ejtm.2023.11856.

[7] C. McDonald et al., “Draft Guidance for Industry Duchenne Muscular Dystrophy, Becker Muscular Dystrophy, and Related Dystrophinopathies – Developing Potential Treatments for the Entire Spectrum of Disease,” JND, vol. 11, no. 2, pp. 499–523, Mar. 2024, doi: 10.3233/JND-230219.

[8] D. J. Birnkrant et al., “Diagnosis and management of Duchenne muscular dystrophy, part 1: diagnosis, and neuromuscular, rehabilitation, endocrine, and gastrointestinal and nutritional management,” Lancet Neurol, vol. 17, no. 3, pp. 251–267, Mar. 2018, doi: 10.1016/S1474-4422(18)30024-3.

[9] E. S. D’Ambrosio and J. R. Mendell, “Evolving Therapeutic Options for the Treatment of Duchenne Muscular Dystrophy,” Neurotherapeutics, vol. 20, no. 6, pp. 1669–1681, Oct. 2023, doi: 10.1007/s13311-023-01423-y.

[10] C. Happi Mbakam and J. P. Tremblay, “Gene therapy for Duchenne muscular dystrophy: an update on the latest clinical developments,” Expert Rev Neurother, vol. 23, no. 10, pp. 905–920, 2023, doi: 10.1080/14737175.2023.2249607.

[11] F. Chemello, E. N. Olson, and R. Bassel-Duby, “CRISPR-Editing Therapy for Duchenne Muscular Dystrophy,” Hum Gene Ther, vol. 34, no. 9–10, pp. 379–387, May 2023, doi: 10.1089/hum.2023.053.

[12] A. Aartsma-Rus, “The Future of Exon Skipping for Duchenne Muscular Dystrophy,” Hum Gene Ther, vol. 34, no. 9–10, pp. 372–378, May 2023, doi: 10.1089/hum.2023.026.

[13] J. M. Spinazzola and L. M. Kunkel, “Pharmacological therapeutics targeting the secondary defects and downstream pathology of Duchenne muscular dystrophy,” Expert Opinion on Orphan Drugs, vol. 4, no. 11, pp. 1179–1194, Nov. 2016, doi: 10.1080/21678707.2016.1240613.

[14] J. Deng, J. Zhang, K. Shi, and Z. Liu, “Drug development progress in duchenne muscular dystrophy,” Front. Pharmacol., vol. 13, p. 950651, Jul. 2022, doi: 10.3389/fphar.2022.950651.

[15] F. Bajanca and L. Vandel, “Epigenetic Regulators Modulate Muscle Damage in Duchenne Muscular Dystrophy Model”.

[16] N. M. Johnson, G. H. Farr, and L. Maves, “The HDAC Inhibitor TSA Ameliorates a Zebrafish Model of Duchenne Muscular Dystrophy,” PLoS Curr, vol. 5, p. ecurrents.md.8273cf41db10e2d15dd3ab827cb4b027, Sep. 2013, doi: 10.1371/currents.md.8273cf41db10e2d15dd3ab827cb4b027.

[17] G. H. Farr et al., “A novel chemical-combination screen in zebrafish identifies epigenetic small molecule candidates for the treatment of Duchenne muscular dystrophy,” Skeletal Muscle, vol. 10, no. 1, p. 29, Oct. 2020, doi: 10.1186/s13395-020-00251-4.

[18] M. Spreafico et al., “Targeting HDAC8 to ameliorate skeletal muscle differentiation in Duchenne muscular dystrophy,” Pharmacological Research, vol. 170, p. 105750, Aug. 2021, doi: 10.1016/j.phrs.2021.105750.

[19] G. C. Minetti et al., “Functional and morphological recovery of dystrophic muscles in mice treated with deacetylase inhibitors,” Nat Med, vol. 12, no. 10, pp. 1147–1150, Oct. 2006, doi: 10.1038/nm1479.

[20] S. Consalvi et al., “Preclinical Studies in the mdx Mouse Model of Duchenne Muscular Dystrophy with the Histone Deacetylase Inhibitor Givinostat,” Mol Med, vol. 19, no. 1, Art. no. 1, Jan. 2013, doi: 10.2119/molmed.2013.00011.

[21] P. Bettica et al., “Histological effects of givinostat in boys with Duchenne muscular dystrophy,” Neuromuscular Disorders, vol. 26, no. 10, pp. 643–649, Oct. 2016, doi: 10.1016/j.nmd.2016.07.002.

[22] E. Mercuri et al., “Safety and efficacy of givinostat in boys with Duchenne muscular dystrophy (EPIDYS): a multicentre, randomised, double-blind, placebo-controlled, phase 3 trial,” The Lancet Neurology, vol. 23, no. 4, pp. 393–403, Apr. 2024, doi: 10.1016/S1474-4422(24)00036-X.

[23] C. Mozzetta, V. Sartorelli, and P. L. Puri, “HDAC inhibitors as pharmacological treatment for Duchenne muscular dystrophy: a discovery journey from bench to patients,” Trends in Molecular Medicine, vol. 30, no. 3, pp. 278–294, Mar. 2024, doi: 10.1016/j.molmed.2024.01.007.

[24] M. Granato et al., “Genes controlling and mediating locomotion behavior of the zebrafish embryo and larva,” Development, vol. 123, no. 1, pp. 399–413, Dec. 1996, doi: 10.1242/dev.123.1.399.

[25] D. I. Bassett, R. J. Bryson-Richardson, D. F. Daggett, P. Gautier, D. G. Keenan, and P. D. Currie, “Dystrophin is required for the formation of stable muscle attachments in the zebrafish embryo,” Development, vol. 130, no. 23, pp. 5851–5860, Dec. 2003, doi: 10.1242/dev.00799.

[26] G. Kawahara and L. M. Kunkel, “Zebrafish based small molecule screens for novel DMD drugs,” Drug Discovery Today: Technologies, vol. 10, no. 1, pp. e91–e96, Mar. 2013, doi: 10.1016/j.ddtec.2012.03.001.

[27] N. B. Wasala, S.-J. Chen, and D. Duan, “Duchenne muscular dystrophy animal models for high-throughput drug discovery and precision medicine,” Expert Opinion on Drug Discovery, vol. 15, no. 4, pp. 443–456, Apr. 2020, doi: 10.1080/17460441.2020.1718100.

[28] J. Berger, S. Berger, T. E. Hall, G. J. Lieschke, and P. D. Currie, “Dystrophin-deficient zebrafish feature aspects of the Duchenne muscular dystrophy pathology,” Neuromuscular Disorders, vol. 20, no. 12, pp. 826–832, Dec. 2010, doi: 10.1016/j.nmd.2010.08.004.

[29] G. Kawahara, J. A. Karpf, J. A. Myers, M. S. Alexander, J. R. Guyon, and L. M. Kunkel, “Drug screening in a zebrafish model of Duchenne muscular dystrophy,” Proceedings of the National Academy of Sciences, vol. 108, no. 13, pp. 5331–5336, Mar. 2011, doi: 10.1073/pnas.1102116108.

[30] M. Karuppasamy et al., “Standardization of zebrafish drug testing parameters for muscle diseases,” Dis Model Mech, vol. 17, no. 1, p. dmm050339, Jan. 2024, doi: 10.1242/dmm.050339.

[31] M. Westerfield, The Zebrafish Book. A Guide for the Laboratory Use of Zebrafish (Danio rerio)., 5th ed. Eugene: university of Oregon Press, 2007.

[32] J. Berger, S. Berger, A. S. Jacoby, S. D. Wilton, and P. D. Currie, “Evaluation of exon-skipping strategies for Duchenne muscular dystrophy utilizing dystrophin-deficient zebrafish,” Journal of Cellular and Molecular Medicine, vol. 15, no. 12, pp. 2643–2651, Dec. 2011, doi: 10.1111/j.1582-4934.2011.01260.x.

[33] D. Aharon and F. L. Marlow, “Sexual determination in zebrafish,” Cell. Mol. Life Sci., vol. 79, no. 1, p. 8, Dec. 2021, doi: 10.1007/s00018-021-04066-4.

[34] E. H. Hasegawa, G. H. Farr, and L. Maves, “Comparison of Pronase versus Manual Dechorionation of Zebrafish Embryos for Small Molecule Treatments,” JDB, vol. 11, no. 2, p. 16, Mar. 2023, doi: 10.3390/jdb11020016.

[35] L. L. Smith, A. H. Beggs, and V. A. Gupta, “Analysis of Skeletal Muscle Defects in Larval Zebrafish by Birefringence and Touch-evoke Escape Response Assays,” JoVE, no. 82, p. 50925, Dec. 2013, doi: 10.3791/50925.

[36] G. Kawahara et al., “Dystrophic muscle improvement in zebrafish via increased heme oxygenase signaling,” Human Molecular Genetics, vol. 23, no. 7, pp. 1869–1878, Apr. 2014, doi: 10.1093/hmg/ddt579.

[37] T. A. Waugh et al., “Fluoxetine prevents dystrophic changes in a zebrafish model of Duchenne muscular dystrophy,” Human Molecular Genetics, vol. 23, no. 17, pp. 4651–4662, Sep. 2014, doi: 10.1093/hmg/ddu185.

[38] Y. B. Kim, K.-H. Lee, K. Sugita, M. Yoshida, and S. Horinouchi, “Oxamflatin is a novel antitumor compound that inhibits mammalian histone deacetylase,” Oncogene, vol. 18, no. 15, pp. 2461–2470, Apr. 1999, doi: 10.1038/sj.onc.1202564.

[39] S. Iezzi, G. Cossu, C. Nervi, V. Sartorelli, and P. L. Puri, “Stage-specific modulation of skeletal myogenesis by inhibitors of nuclear deacetylases,” Proc Natl Acad Sci U S A, vol. 99, no. 11, pp. 7757–7762, May 2002, doi: 10.1073/pnas.112218599.

[40] S. Iezzi et al., “Deacetylase inhibitors increase muscle cell size by promoting myoblast recruitment and fusion through induction of follistatin,” Dev Cell, vol. 6, no. 5, pp. 673–684, May 2004, doi: 10.1016/s1534-5807(04)00107-8.

[41] S. Consalvi et al., “Determinants of epigenetic resistance to HDAC inhibitors in dystrophic fibro-adipogenic progenitors,” EMBO Rep, vol. 23, no. 6, p. e54721, Jun. 2022, doi: 10.15252/embr.202254721.

[42] J. J. Widrick, G. Kawahara, M. S. Alexander, A. H. Beggs, and L. M. Kunkel, “Discovery of Novel Therapeutics for Muscular Dystrophies using Zebrafish Phenotypic Screens,” J Neuromuscul Dis, vol. 6, no. 3, pp. 271–287, 2019, doi: 10.3233/JND-190389.

[43] E. E. Patton, L. I. Zon, and D. M. Langenau, “Zebrafish disease models in drug discovery: from preclinical modelling to clinical trials,” Nat Rev Drug Discov, vol. 20, no. 8, pp. 611–628, Aug. 2021, doi: 10.1038/s41573-021-00210-8.

[44] H.-C. Lee, C.-Y. Lin, and H.-J. Tsai, “Zebrafish, an In Vivo Platform to Screen Drugs and Proteins for Biomedical Use,” Pharmaceuticals (Basel), vol. 14, no. 6, p. 500, May 2021, doi: 10.3390/ph14060500.

[45] N. Ohnesorge, T. Sasore, D. Hillary, Y. Alvarez, M. Carey, and B. N. Kennedy, “Orthogonal Drug Pooling Enhances Phenotype-Based Discovery of Ocular Antiangiogenic Drugs in Zebrafish Larvae,” Front Pharmacol, vol. 10, p. 508, 2019, doi: 10.3389/fphar.2019.00508.

[46] C. Mozzetta et al., “Fibroadipogenic progenitors mediate the ability of HDAC inhibitors to promote regeneration in dystrophic muscles of young, but not old Mdx mice,” EMBO Molecular Medicine, vol. 5, no. 4, pp. 626–639, Apr. 2013, doi: 10.1002/emmm.201202096.

[47] V. Saccone et al., “HDAC-regulated myomiRs control BAF60 variant exchange and direct the functional phenotype of fibro-adipogenic progenitors in dystrophic muscles,” Genes Dev., vol. 28, no. 8, pp. 841–857, Apr. 2014, doi: 10.1101/gad.234468.113.

[48] M. Sandonà et al., “Histone Deacetylases: Molecular Mechanisms and Therapeutic Implications for Muscular Dystrophies,” IJMS, vol. 24, no. 5, p. 4306, Feb. 2023, doi: 10.3390/ijms24054306.

[49] C. Colussi et al., “HDAC2 blockade by nitric oxide and histone deacetylase inhibitors reveals a common target in Duchenne muscular dystrophy treatment,” Proceedings of the National Academy of Sciences, vol. 105, no. 49, pp. 19183–19187, Dec. 2008, doi: 10.1073/pnas.0805514105.

[50] A. Zutz, L. Chen, F. Sippl, A. Humpe, and C. Schölz, “Epigenetic Compound Screening Uncovers Small Molecules for Reactivation of Latent HIV-1,” Antimicrob Agents Chemother, vol. 65, no. 1, pp. e01815–20, Dec. 2020, doi: 10.1128/AAC.01815-20.

[51] A. Osseni et al., “Pharmacological inhibition of HDAC6 improves muscle phenotypes in dystrophin-deficient mice by downregulating TGF-β via Smad3 acetylation,” Nat Commun, vol. 13, no. 1, p. 7108, Nov. 2022, doi: 10.1038/s41467-022-34831-3.

[52] N. Erdeljac, K. Bussmann, A. Schöler, F. K. Hansen, and R. Gilmour, “Fluorinated Analogues of the Histone Deacetylase Inhibitor Vorinostat (Zolinza): Validation of a Chiral Hybrid Bioisostere, BITE,” ACS Med Chem Lett, vol. 10, no. 9, pp. 1336–1340, Sep. 2019, doi: 10.1021/acsmedchemlett.9b00287.

[53] B. Zhang, J. Liu, D. Gao, X. Yu, J. Wang, and X. Lei, “A fluorine scan on the Zn2+-binding thiolate side chain of HDAC inhibitor largazole: Synthesis, biological evaluation, and molecular modeling,” Eur J Med Chem, vol. 182, p. 111672, Nov. 2019, doi: 10.1016/j.ejmech.2019.111672.

